# Mixed DAMP/MAMP oligosaccharides promote both growth and defense against fungal pathogens of cucumber

**DOI:** 10.1101/2024.12.27.630494

**Authors:** Sreynich Pring, Hiroaki Kato, Keiko Taniuchi, Maurizio Camagna, Makoto Saito, Aiko Tanaka, Bryn A. Merritt, Orlando Argüello-Miranda, Ikuo Sato, Sotaro Chiba, Daigo Takemoto

## Abstract

Plants recognize a variety of environmental molecules, thereby triggering appropriate responses to biotic or abiotic stresses. Substances containing microbes-associated molecular patterns (MAMPs) and damage-associated molecular patterns (DAMPs) are representative inducers of pathogen resistance and damage repair, thus treatment of healthy plants with such substances can pre-activate plant immunity and cell repair functions. In this study, the effects of DAMP/MAMP oligosaccharides mixture (Oligo-Mix) derived from plant cell wall (cello-oligosaccharide and xylo-oligosaccharide), and fungal cell wall (chitin-oligosaccharide) were examined in cucumber. Treatment of cucumber with Oligo-Mix promoted root germination and plant growth, along with increased chlorophyll contents in the leaves. Oligo-Mix treatment also induced typical defense responses such as MAP kinase activation and callose deposition in leaves. Pretreatment of Oligo-Mix enhanced disease resistance of cucumber leaves against pathogenic fungi *Podosphaera xanthii* (powdery mildew) and *Colletotrichum orbiculare* (anthracnose). Oligo-Mix treatment increased the induction of hypersensitive cell death around the infection site of pathogens, which inhibited further infection and the conidial formation of pathogens on the cucumber leaves. RNA-seq analysis revealed that Oligo-Mix treatment upregulated genes associated with plant structural reinforcement, responses to abiotic stresses and plant defense. These results suggested that Oligo-Mix has beneficial effects on growth and disease resistance in cucumber, making it a promising biostimulant for agricultural application.

## 1. Introduction

Plants have an array of mechanisms for sensing and adapting to environmental stresses. Some breakdown molecules of plant cells are released by physical damage, pathogen infection, or insect attack and are recognized as signals by plants to promote cellular repair or defense responses (Tanaka and Heil 2021; Ge et al., 2022; Chen et al., 2024). Such molecules are commonly called DAMPs (damaged-associated molecular patterns), including cello-oligosaccharides (COS), which are constituent molecules of cellulose derived from plant cell walls, and xylo-oligosaccharides (XOS) contained in hemicellulose, which cross-link cellulose and lignin in the cell wall (Souza et al., 2017; Claverie et al., 2018; Pring et al., 2023). Volatiles emitted by damaged plants are also involved in stress responses, thus considered as a type of DAMPs (Yamauchi et al., 2018). It is also known that specific peptides released externally by cell disruption are recognized as DAMPs by receptors on the surface of neighboring cells (Pearce et al., 1991; Yamaguchi et al., 2010; Wang et al., 2018; Hander et al., 2019).

Molecules specific to microorganisms are recognized by plant cells as alert signals of pathogen attack, and the epitope structures of such substances are called MAMPs (microbes-associated molecular patterns, Ge et al., 2022). Some MAMPs are highly conserved in large groups (i.e. bacteria, fungi or oomycetes), such as flg22 (a conserved peptide of the bacterial flagellar protein flagellin, Gómez-Gómez et al., 1999), peptidoglycan (the structural polymer of the bacterial cell wall, Gust et al., 2007), chitin and β-glucan (major cell wall components of fungi, Sharp and Albersheim, 1984; Felix et al., 1993) and 9-methyl-4,8-sphingadienine (9Me-Spd, a substructure of ceramides found in the cell membrane of fungi and oomycetes, Kato et al., 2022; Monjil et al., 2024). Other MAMPs are found in certain groups of pathogenic microbes, such as elicitins produced by oomycete pathogens (Derevnina et al., 2016), and fungal ethylene-inducing xylanases (EIXs, Dean et al., 1991). On the other hand, the recognition mechanisms of MAMPs by plant species are also diverse; for example, Solanaceae plants and rice can recognize different parts of bacterial flagellin other than flg22 (Fliegmann et al., 2016). Elicitin and EIX have also been reported to be recognized as MAMPs by a limited range of plant species (Kamoun et al., 1993; Ron and Avni, 2004; Takemoto et al., 2005).

While the understanding of the recognition mechanisms of DAMPs and MAMPs by plants is a subject of basic research, these substances are also anticipated to be utilized as materials to activate plant growth and immunity in agricultural production. Substances that activate beneficial plant responses are collectively referred to as biostimulants. The DAMPs and MAMPs derived from fundamental structures of plants and microbes are expected to be versatile as agricultural materials because they can activate the responses of diverse plant species. The cell walls of plants and fungi are essentially composed of common materials (Cosgrove 2005; Gow and Lenardon 2023), thus substances in their cell walls are representative candidates for biostimulant.

Previously, we have shown that DAMP and MAMP oligosaccharides, COS prepared from cotton linters, XOS prepared from corn cobs as well as chitin-oligosaccharide (CHOS) from crustacean shells can trigger typical defense responses of Arabidopsis such as production of reactive oxygen species (ROS), MAP kinases phosphorylation and callose depositions (Pring et al., 2023). Gene ontology enrichment analysis of RNA-seq data revealed that simultaneous treatment of COS, XOS and CHOS (Oligo-Mix) effectively activate genes for disease resistance. Generally, it is considered that activating disease resistance in plants suppresses their growth, referred to as “trade-off effect” (Karasov et al., 2017; He et al., 2022; Godínez-Mendoza et al., 2023; Gao et al., 2024). However, the treatment of Oligo-Mix had no significant defects on the expression of photosynthesis genes. In practice, treatment of the Oligo-Mix enhanced resistance of tomato against powdery mildew pathogen, and moreover, tomato growth was rather promoted by Oligo-Mix treatment (Pring et al., 2023).

In this study, the effects of Oligo-Mix treatment on cucumber were investigated. Cucumber (*Cucumis sativus* L.) is one of the most important vegetable crops cultivated worldwide. Besides the varieties with higher resistance to pathogens, in general, cucumbers are susceptible to a variety of destructive pathogens. Anthracnose, caused by a fungal pathogen *Colletotrichum orbiculare*, is a major foliage disease that reduces the yield and quality of cucumber and other Cucurbitaceae plants such as watermelon and melon. *C. orbiculare* is a hemi-biotrophic pathogen that employs both biotrophic and necrotrophic phases to infect its hosts (Matsuo et al., 2022). Powdery mildew of cucumber is widely distributed, rapidly spreading to cause destructive disease by obligate parasites *Golovinomyces cichoracearum* or *Podosphaera xanthii* (Lebeda et al., 2011). While there are some fungicides accessible for managing these pathogens, the emergence of drug-resistant fungal strains has become a major problem (Vielba-Fernández et al., 2020). Here, we evaluated the influence of the Oligo-Mix treatment on cucumber growth, development, and resistance against anthracnose and powdery mildew pathogens.

## 2. Material and methods

### 2.1. Plant material and growth condition

Cucumber (cv. Shimoshirazu Jibai, Takii seed, Kyoto, Japan) were grown in autoclaved commercial soil (Sakata Super Mix A, Sakata seed, Yokohama, Japan) in a plant growth room at 23°C with 16 h of light per day. For the greenhouse test, two-week-old seedlings grown in a controlled growth chamber were transferred to pots (28 cm diameter x 22 cm height, filled with Sakata Super Mix A) and grown in a greenhouse in Togo field, Nagoya University (University farm, Togo, Aichi, Japan).

### 2.2. Oligosaccharide mixture (Oligo-Mix)

Oligo-Mix and control solutions were provided by Resonac Corporation (Tokyo, Japan). The purity and average degrees of polymerization (DP) of oligosaccharides used in this study were confirmed by GPC (gel permeation chromatography). Oligo-Mix (undiluted solution) used in this study consists of 20 mg/ml cello-oligosaccharide (COS, prepared from cotton linter, purity >99%, average DP = 3.4), 40 mg/ml xylo-oligosaccharide (XOS, from corn cob, purity >99%, average DP = 2.7), and 20 mg/ml chitin-oligosaccharide (CHOS, from shrimp shell, purity >99%, average DP = 3.5) (Sreynich et al., 2023). Both Oligo-Mix and the control solutions contain water-soluble P (P_2_O_5_, 4.7% w/v) and K (K_2_O, 3.4% w/v). Oligo-Mix and control solution were diluted to 1/1000 and used for the treatment of plants by spray or infiltration using a needleless syringe.

### 2.3. Determination of root number and length

Cucumber seeds were placed on laboratory tissue soaked with distilled water in Petri dishes and incubated at 26 °C with a light/dark cycle set at 16 h/8 h. The cucumber seeds germinated at same day were collected and sprayed with Oligo-Mix or control solution (Pressed the sprayer 3 times, approx. 250 µl per petri dish), and the number of roots per seed and the length of longest root were measured 2 days after the treatment.

### 2.4. Determination of cucumber growth

The individual seedlings, treated with control solution or Oligo-Mix as described above, were transferred into 9-cm-diameter plastic pot filled with sterilized soil mix and grown in growth room at 23 °C with 16 h of light per day. Cucumber plants were sprayed with Oligo-Mix or control solution once every week (Pressed the sprayer 24 times, approx. 2 ml per plant, 4 times altogether) until sampling. After one week of the last treatment, the fresh weight of the root and above-ground tissues are measured.

### 2.5. Measurement of chlorophyll contents

Chlorophyll was extracted according to the method described in Shibata et al., (2016). One g of second true leaves of cucumber was soaked in 50 ml methanol overnight. The absorbance intensity of the solution was quantified by spectrophotometer (Multiskan GO, Thermo Fisher Scientific, Waltham, MA, USA), and the concentration of chlorophyll a and b was calculated according to the method of Porra and Scheer (1989) as follows; Chlorophyll a (μg/ml) = 16.29 x A^665^ − 8.54 x A^652^; Chlorophyll b (μg/ml) = 30.66 x A^652^ − 13.58 x A^665^.

### 2.6. MAP kinase activation assay

Activation of MAP kinases in Arabidopsis was detected as previously reported (Kato et al., 2022). Cucumber seedlings were treated with Oligo-Mix or control solution (by infiltration with a needleless syringe) for 15 or 30 min and frozen in liquid nitrogen. Proteins were extracted in extraction buffer (50 mM HEPES-KOH pH 7.4, 5 mM EDTA, 0.5 mM EGTA, 50 mM ß-glycerophosphate, 10 mM NaF, 10 mM Na_3_VO_4_, and 2 mM DTT). Phosphorylated MAP kinases were detected by western blot using Phospho-p44/42 MAPK (Erk1/2; Thr-202/Tyr-204) rabbit monoclonal antibodies #9101 (Cell Signaling Technology). Blots were stained with PageBlue Protein Staining Solution (Thermo Fisher Scientific) to verify equal loading.

### 2.7. Staining of callose deposition

Staining of callose deposition was performed as previously reported (Shibata et al., 2016). Cucumber seedlings were treated with Oligo-Mix or control solution by infiltration with needleless syringe, and seedlings were fixed 24 h after treatment in fixation solution (1% [v/v] glutaraldehyde, 5 mM citric acid, and 90 mM Na_2_HPO_4_, pH 7.4) overnight. Fixed seedlings were heated in boiled water for 5 minutes, decolorized in ethanol, and stained in aniline blue stain (0.1% [w/v] aniline blue in 67 mM phosphate buffer, pH 12.0) to detect callose deposition. The fluorescence spots of deposited callose were detected by a fluorescence microscope (BX51, Olympus, Tokyo, Japan) using an excitation wavelength of 365 nm. The number of callose spots per area was quantified using ImageJ software (Schneider et al., 2012).

### 2.8. Pathogens and inoculation

Powdery mildew of cucumber, *Podosphaera xanthii* strain SP23, was isolated from cucumber grown in a greenhouse in Togo field, Nagoya University, and maintained by inoculating healthy cucumbers every 5-10 days. The species of powdery mildew was determined based on the sequencing of ITS (Internal transcribed spacer) region using the method previously described (Ashida et al., 2023). Conidia newly formed on cucumber leaves were suspended in water and adjusted to a concentration of 1×10^5^ spores/ml. The 5-leaf-stage cucumber plants were inoculated by spraying 0.5 ml spore suspension per plant at 24 h after the treatment with control solution or Oligo-Mix. One week later, spot symptoms were observed on the leaf surface, and number and type (spored, yellowish, or cell death) of colony were counted. To measure conidial formation, cut leaves with colonies (1.5 × 1.5 cm) were scrubbed with 2 ml of distilled water using a cotton bud, then filtered with a stainless filter disc (mesh size 100 µm). Subsequently, the number of conidia was quantified with a hemocytometer (Erma, Tokyo, Japan).

Anthracnose pathogen, *Colletotrichum orbiculare* strain T-104 (MAFF240422, which was isolated by Dr. Yasumori, Kyoto University in 1951) was provided by Prof. Yoshitaka Takano (Kyoto University, Japan). *C. orbiculare* was grown on potato dextrose agar (PDA) in 23°C for 7 days, and conidia formed on the colony were suspended in water and adjusted to a concentration of 1×10^5^ spores/ml for the inoculation. Cucumber was spray-inoculated with 0.5 ml of spore suspension per plant. The symptoms observed on the leave were photographed 7 days after the inoculation.

### 2.9. Detection of anthracnose lesions on cucumber leaves

Briefly, the area affected by anthracnose on cucumber leaves was calculated as the percentage of the affected area divided by the total leaf area. The calculation of the total leave area was done using the custom-made MATLAB script (OAM_240209_leave_seg) to isolate all pixels in an image that belonged to a leaf according to a median intensity threshold. To identify specific infected areas on individual leaves, we trained a modified pixel flow-driven CNN architecture (Stringer et al., 2021) to identify each pixel on a leaf that could be classified as affected with more than 0.9 probability. The CNN model was trained using the preliminary segmentations from the MATLAB script based on median thresholding, which generated single-leaf masks depicting the affected areas. The preliminary masks were manually corrected using custom-made software to remove incorrectly classified pixels, and the final corrected images were split into training (70%), and testing (30%) data sets for deep learning model training. The CNN architecture was modified to obtain a binary lesion/no-lesion single-pixel classification (semantic segmentation) as direct output. The CNN model “Takemoto_1” and the accessory code for whole leaf segmentation, and to obtain the final semantic segmentation from the CNN, as well as the original image dataset with labels and a toy dataset to test the segmentation code, are freely available from the GitHub repository of the Miranda Lab: https://github.com/MirandaLab/Deep_Leaf_Segmentation. Leaf segmentations were performed on a Dual Intel Xeon Silver 4216 (2.1GHz,3.2GHz Turbo, 16C,9.6GT/s 2UPI, 22MB Cache, HT (100W) equipped with an Nvidia Quadro RTX5000, 16GB, 4DP, VirtualLink (XX20T) outfitted with 64GB 4×16GB DDR4 2933MHz RDIMM ECC Memory and a 2TB primary NVMe SSD. Differences between control and treatments were evaluated using the Kolmogorov-Smirnov tests *kstest2* MATLAB function, with significance set at p < 0.05.

### 2.10. Trypan blue staining and microscopy

To visualize the pathogen hyphae and conidiophores, or HR-like cell death of plant cells, the infected leaves were stained with trypan blue as described previously (Takemoto et al., 2003). Small pieces of leaves were cleared in methanol in 1.5 ml tube for more than 24 h (methanol was changed once) until the chlorophyll (green color) was removed. Then methanol was replaced with lactophenol trypan blue stain (10 ml lactic acid, 10 ml glycerol, 10 g phenol, 10 mg trypan blue, 10 ml H_2_O) and heated in boiled water for 3 mins. After cooling down to room temperature, the stain was replaced with 1 g/ml chloral hydrate, and gently shaken overnight. Pathogen hyphae and plant cells were monitored under a light microscope (BX51, Olympus, Japan).

### 2.11. RNA-seq and data analysis

The four-weeks old cucumber plants were sprayed with Oligo-Mix or control solution and total RNA was isolated at 24 h after treatment. RNA extraction was performed using the RNeasy Plant Mini Kit (QIAGEN, Hilden, Germany). Evaluation of RNA quality, construction of library and sequencing were performed principally as previously described (Kuroyanagi et al., 2022, Bulasag et al., 2023). Libraries were constructed using KAPA mRNA Capture Kit (Roche Diagnostics, Tokyo, Japan) and MGIEasy RNA Directional Library Prep Set (MGI, Shenzhen, China), and sequenced on DNBSEQ-G400RS (MGI) with 150 bp paired-end protocol. The RNA-seq reads were filtered using trim-galore v.0.6.6 (Martin, 2011, bioinformatics.babraham.ac.uk) and mapped to the cucumber genome (genome assembly Cucumber_9930_V3, NCBI RefSeq sequence GCA_000004075.3) using HISAT2 v.2.2.1 (Kim et al., 2019) and abundance inferred via StringTie v.2.1.7 (Kovaka et al., 2019). Significant differential expression was determined using DESeq2 v.1.32.0 (Love et al., 2014). All software used during RNA-seq analysis was run with default settings. RNA-seq data for cucumber reported in this work are available in GenBank under the accession numbers DRA019928. Gene ontology (GO) enrichment analysis was performed using Arabidopsis homologue of cucumber genes upregulated by Oligo-Mix treatment (Table S1). PANTHER statistical overrepresentation test (http://pantherdb.org; Version 17.0, Thomas et al. 2022) using default settings (Fisher’s exact test, False discovery rate (FDR) < 0.05). SRplot (https://www.bioinformatics.com.cn/en), a free online data analysis and visualization platform, was used to draw the graphs.

## 3. Results and discussion

### 3.1. Effect of Oligo-Mix treatment on the growth of cucumber

To elucidate the effect of the Oligo-Mix (mixture of cello-oligosaccharide COS, xylo-oligosaccharide XOS, and chitin-oligosaccharide CHOS) on cucumber growth, newly germinated cucumber seeds were spray-treated with Oligo-Mix or control solution, and root number and length were measured 2 days after the treatment. The number of roots per seed and length of longest root were significantly increased (almost 2 times), indicating that Oligo-Mix can promote the growth of cucumber roots (Fig. 1 A). Those seedlings were transplanted to the pot and sprayed with Oligo-Mix weekly (once a week). One week after the last spray (sprayed 4 times and 5 weeks after transplant), the growth of cucumber plants was evaluated. The fresh weight of roots and above-ground tissue was significantly increased by Oligo-Mix treatment (Fig. 1 B). Previously, we have reported the positive effects of Oligo-Mix on root and plant growth of tomato (Sreynich et al., 2023). Additionally, leaves of cucumber treated with Oligo-Mix exhibited a visibly darker green compared to those in the control group. To evaluate the effect of Oligo-Mix on chlorophyll contents, cucumber leaves were immersed with methanol to extract and calculated chlorophylls according to the method reported by Porra and Scheer (1989). Chlorophyll a in leaves treated with Oligo-Mix was approx. 405 µg/ml, whereas control leaves contained 256 µg/ml. Similarly, chlorophyll b contents was 143 µg/ml in leaves treated with Oligo-Mix, while at 89 µg/ml for control leaves. These results suggest that Oligo-Mix can effectively enhance photosynthetic capacity, probably leading to improved plant growth.

**Fig. 1.**
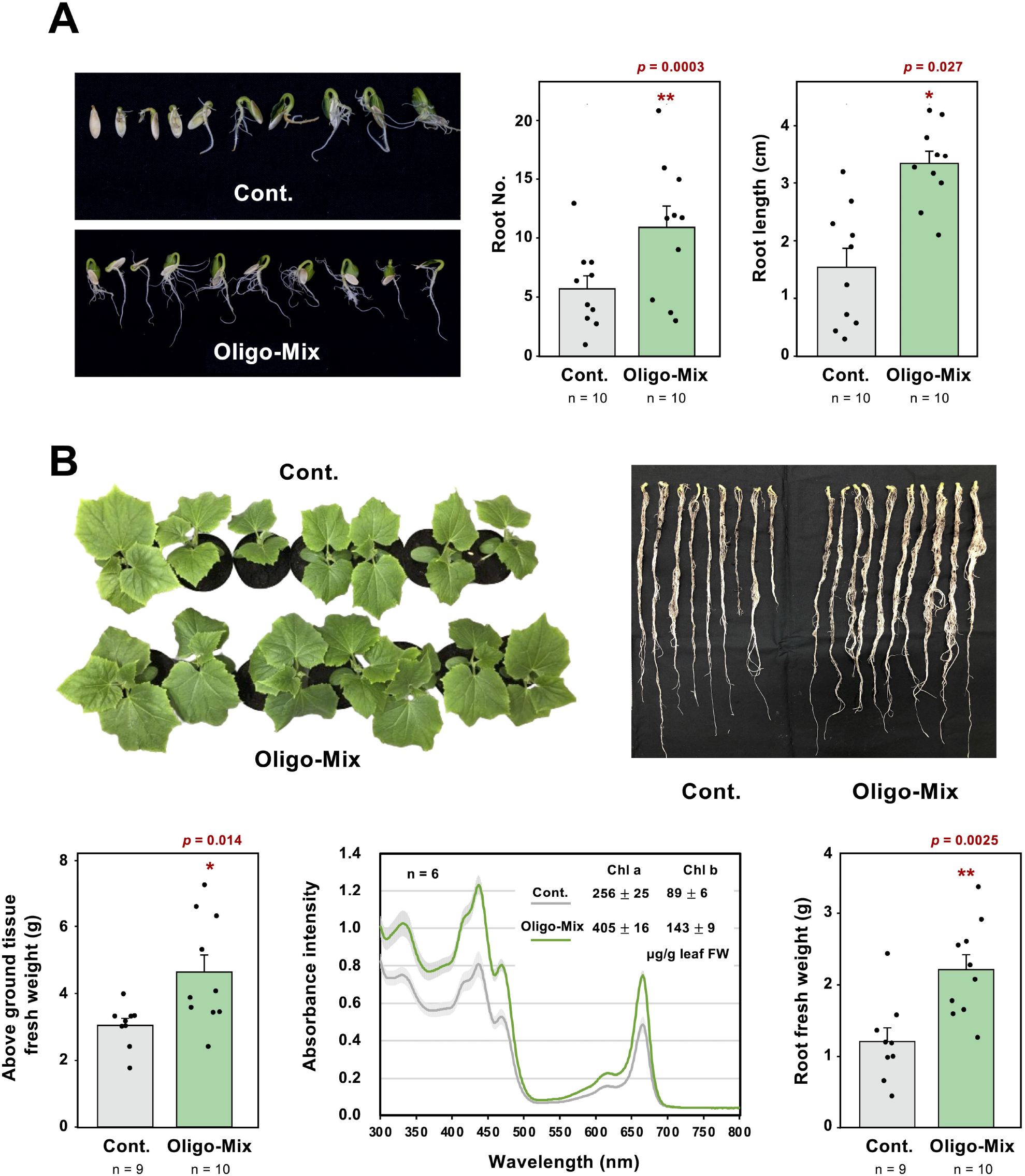
Oligo-Mix treatment can promote the growth of cucumber. **(A)** Effect of Oligo-Mix treatment on seed germination of cucumber. Cucumber seeds were placed on laboratory tissue soaked with distilled water in Petri dishes, and germinated seeds were sprayed with control solution or Oligo-Mix (containing 20 µg/ml cello-oligosaccharide [COS], 40 µgl/m1 xylo-oligosaccharide [XOS], and 20 µg/ml chitin-oligosaccharide [CHOS]). The number of roots and root length (longest root) were measured 2 days after spray. **(B)** Effect of Oligo-Mix treatment on growth of cucumber. Cucumber plants (28 days old) were sprayed with control solution or Oligo-Mix every week (4 times), and fresh weight of above-ground tissue (left) and root (right) were measured 7 days after the last treatment. (middle) Chlorophyll contents in cucumber leaves were quantified according to the method described in Porra and Scheer (1989). Asterisks indicate a significant difference from control as assessed by a two-tailed Student’s *t*-test. * p < 0.05, ** p < 0.01.

### 3.2 Induction of MAP kinase phosphorylation and callose deposition of cucumber by Oligo-Mix treatment

Activation of MAP kinase cascade is a common initial response of plants to induce disease resistance (Asai et al., 2002; Sun and Zhang, 2022). Cucumber cotyledons or Arabidopsis seedlings were treated with Oligo-Mix and phosphorylated MAP kinases were detected using antibody against phospho-MAP kinases (Fig. 2A). Within 15 min after Oligo-Mix treatment, rapid activation of cucumber and Arabidopsis MAP kinases was detected. In Arabidopsis, AtMPK6 (approx. 44 kDa) is the major MAP kinase activated by Oligo-Mix, and relatively minor activation of AtMPK3 and AtMPK4 (approx. 39 kDa and 37 kDa, respectively) were also detected. Similarly, in cucumber, a major activated MAP kinase (approx. 46 kDa) and weak activation of two MAP kinases were also detected within 15 min after Oligo-Mix treatment, suggesting that analogous activation of similar MAP kinases is induced by Oligo-Mix treatment in Arabidopsis and cucumber.

**Fig. 2.**
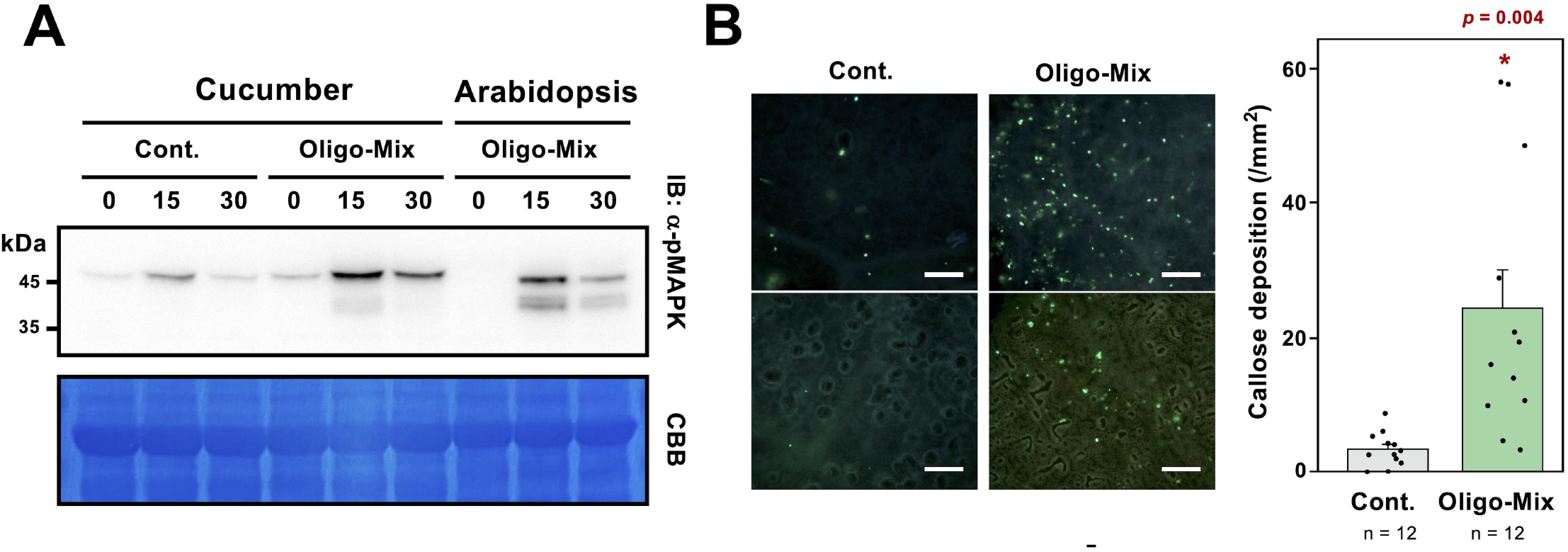
Oligo-Mix treatment induces activation of MAP kinase and callose deposition of cucumber. **(A)** Cucumber or Arabidopsis seedlings were treated with control solution (Cont.) or Oligo-Mix (containing 20 µg/ml cello-oligosaccharide, 40 µg/m1 xylo-oligosaccharide and 20 µg/ml chitin-oligosaccharide). The phosphorylation of MAP kinases was determined by western blot using phospho-p44/42 MAPK (Erk1/2; Thr-202/Tyr-204) antibody. **(B)** Cucumber seedlings were treated with control solution (Cont.) or Oligo-Mix and callose deposition was visualized by aniline blue staining. The fluorescence spots were counted 24 h after treatment under a fluorescence microscope (n = 12). Bar = 200 μm.

Callose deposition is another typical plant defense response to reinforce plant cell walls during pathogen attack (Ellinger and Voigt, 2014). After treatment of cucumber cotyledons with Oligo-Mix for 24 h, a significant increase in callose depositions was detected (Fig. 2B). These results suggested that Oligo-Mix treatment can promote the defense responses of cucumber.

### 3.3 *Effect of Oligo-Mix* treatment on disease resistance of cucumber against powdery mildew and anthracnose pathogens

Two types of plant pathogens were employed to examine the effect of Oligo-Mix on plant disease resistance. Cucumber powdery mildew fungus, *Podosphaera xanthii*, is an obligate parasitic pathogen of Cucurbitaceae. Cucumber plants were sprayed with control solution or Oligo-Mix, and inoculated with spore suspension of *P. xanthii* 24 h after the treatment. Although the number of lesions developed on leaves was not affected by Oligo-Mix treatment, the ratio of yellowish or necrotic lesions was significantly increased for leaves treated with Oligo-Mix (Fig. 3). Microscopic observation revealed the successful penetration, formation of haustoria, and massive production of conidiophores and conidia on control leaves, while HR (hypersensitive response)-like cell death was often found for the leaves treated with Oligo-Mix (Fig. 4A). Consistently, Oligo-Mix treatment significantly reduced the spore formation on the leaves (Fig. 4B and C). These results indicate that the Oligo-Mix treatment is effective in reducing the amount of spore formation of powdery mildew, thus presumed to be effective in suppressing the secondary spread of the disease.

**Fig. 3.**
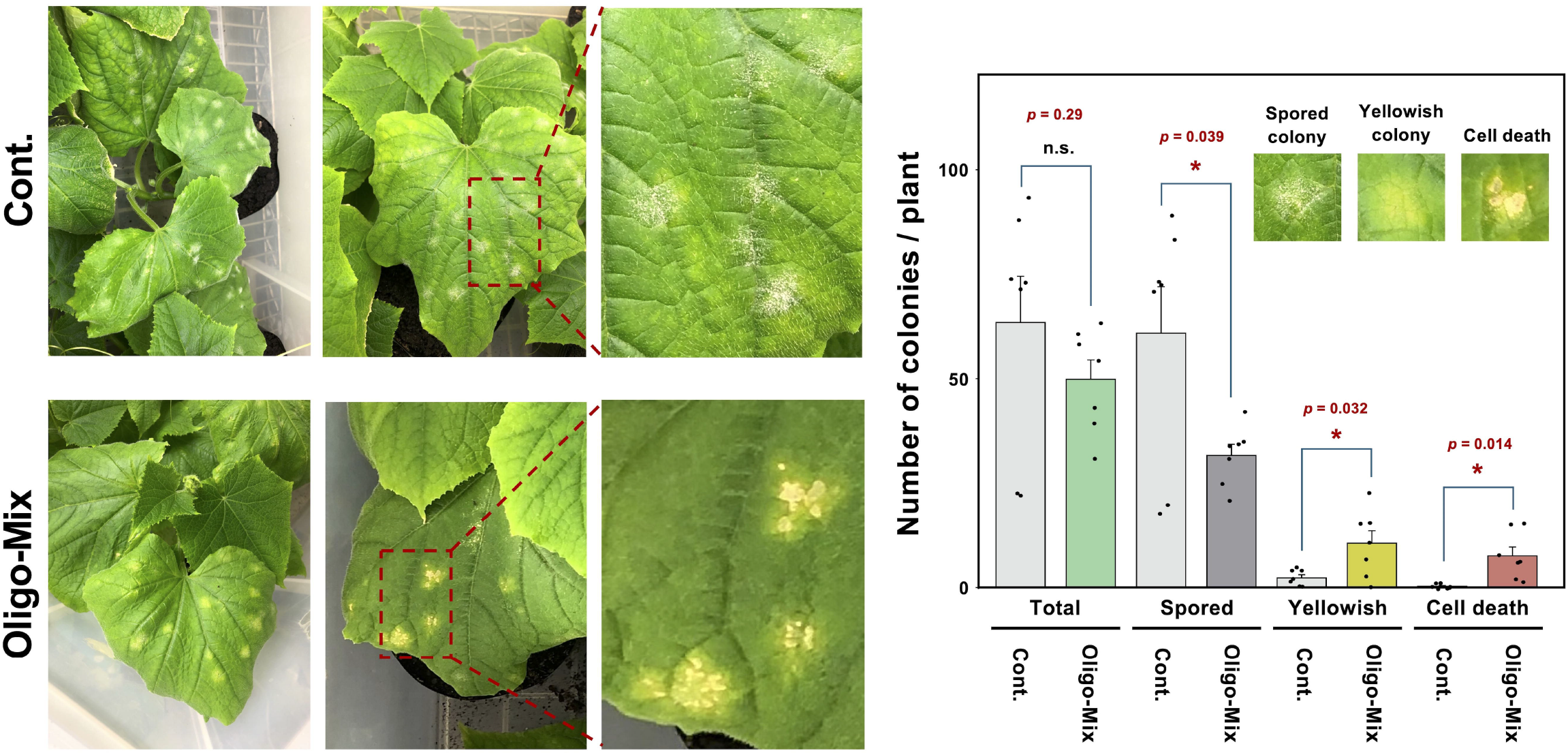
Oligo-Mix induces disease resistance against powdery mildew (*Podosphaera xanthii*) on cucumber. Cucumber plants (4-weeks-old) were treated with control solution (Cont.) or Oligo-Mix (containing 20 µg/ml cello-oligosaccharide, 40 µg/m1 xylo-oligosaccharide and 20 µg/ml chitin-oligosaccharide) and spray-inoculated with powdery mildew spore suspension (1×10^5^ spores/ml, 0.5 ml/plant) at 24 h after treatment. The number of powdery mildew colony types (spored, yellowish, or cell death) per plant was counted one week after the inoculation (n = 7). Asterisks indicate a significant difference from the control as assessed by a two-tailed Student’s *t*-test. * *p* < 0.05. n.s., not significant.

**Fig. 4.**
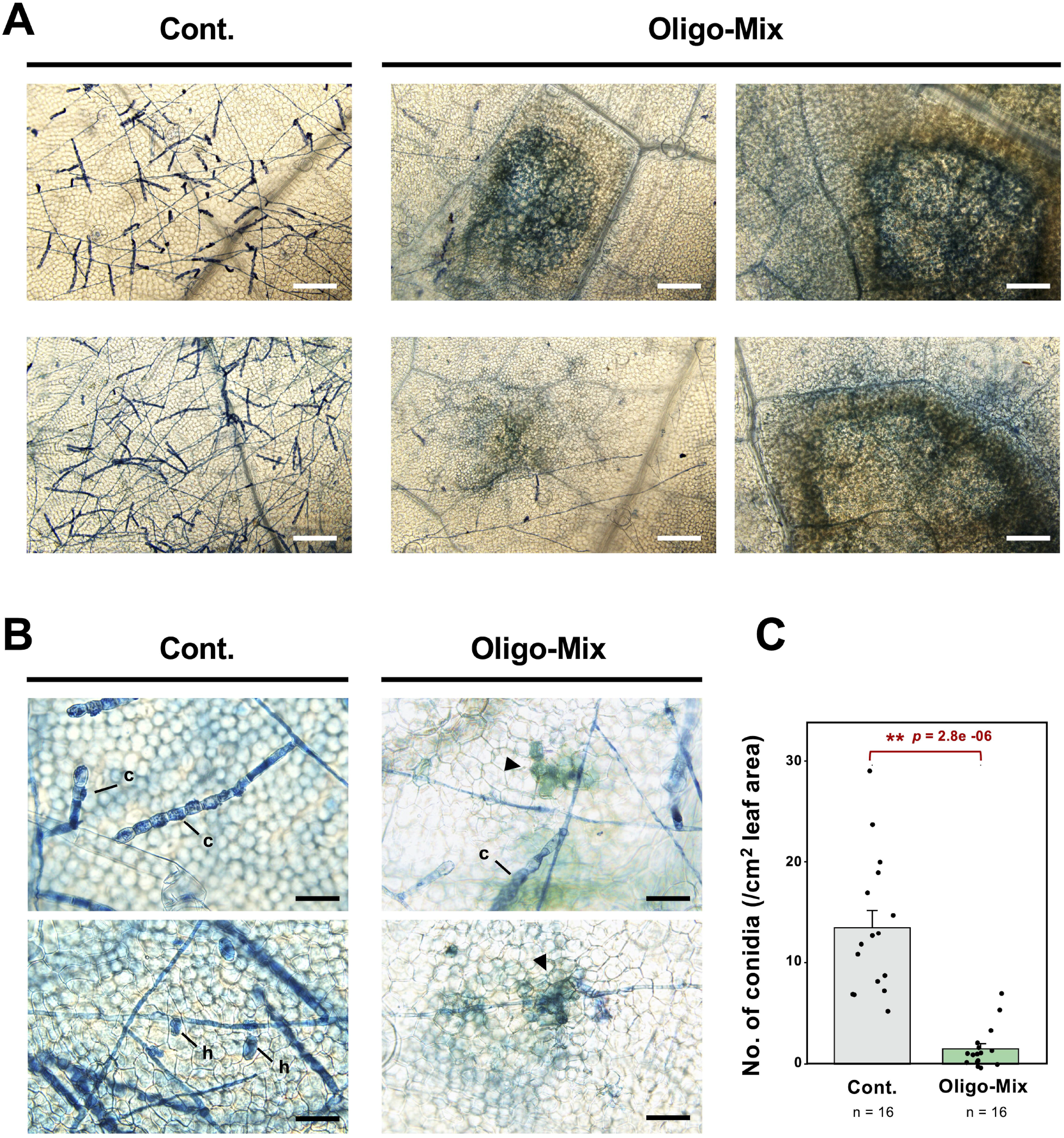
Pretreatment of Oligo-Mix can promote the resistance of cucumber against powdery mildew pathogen *Podosphaera xanthii*. Cucumber plants (4-weeks-old) were treated with control solution (Cont.) or Oligo-Mix (containing 20 µg/ml cello-oligosaccharide, 40 µg/m1 xylo-oligosaccharide and 20 µg/ml chitin-oligosaccharide) and spray-inoculated with powdery mildew spore suspension (1×10^5^ spores/ml, 0.5 ml/plant) at 24 h after treatment. Inoculated leaves were stained with lacto-phenol trypan blue 8 days after inoculation. **(A)** Extensive hyphal growth and conidiophores formation were observed on cucumber leaves 8 days after the inoculation, while induction of cell death was detected for the leaves pretreated with Oligo-Mix. Bars = 200 µm. **(B)** Fully developed conidiophores (c) and haustoria (h) were observed on the leaves of control plants. Local cell death (arrowheads) at the sites of penetration/haustoria formation of *P. xanthii* was detected on cucumber leaves pretreated with Oligo-Mix. Bars = 50 µm. **(C)** Number of conidia formed on leaves (with symptoms) was counted 7 days after the inoculation (n = 16). Asterisks indicate a significant difference from the control as assessed by a two-tailed Student’s *t*-test. ** *p* < 0.01.

*Colletotrichum orbiculare*, a cucurbit anthracnose fungus, is a hemibiotrophic pathogen that infects cucumber, melon, watermelon, and other cucurbitaceous plants. As the experiment for the powdery mildew, cucumber leaves treated with control solution or Oligo-Mix were inoculated with spore suspension of *C. orbiculare* 24 h after the treatment. Yellow primary lesions appeared on inoculated leaves about 4 days after inoculation and were obvious by 7 days. Image analysis to quantify the percentage of infected and yellowed areas (see details in Material and methods section) indicated that Oligo-Mix treatment reduced the severity of the anthracnose disease (Fig. 5). In control leaves, necrotic lesions and outgrowth of intracellular biotrophic hyphae from necrotic area were observed (Fig. 6). Formation of acervuli and setae (asexual fruiting body), and extensive growth of hyphae on leaf surface were also observed in control leaves, indicating the successful hemibiotrophic infection of anthracnose pathogen. For leaves treated with Oligo-Mix, in contrast, only immature acervuli and setae were found and necrotic lesions with no further extension of pathogen hyphae were observed (Fig. 6). These observations indicated that pre-treatment of Oligo-Mix can reduce the infection of anthracnose pathogen. These results indicate that pretreatment with Oligo-Mix is effective against biotrophic and hemibiotrophic pathogens. However, it should be noted that its effect for plant resistance against necrotrophic pathogens is unknown and requires further investigation.

**Fig. 5.**
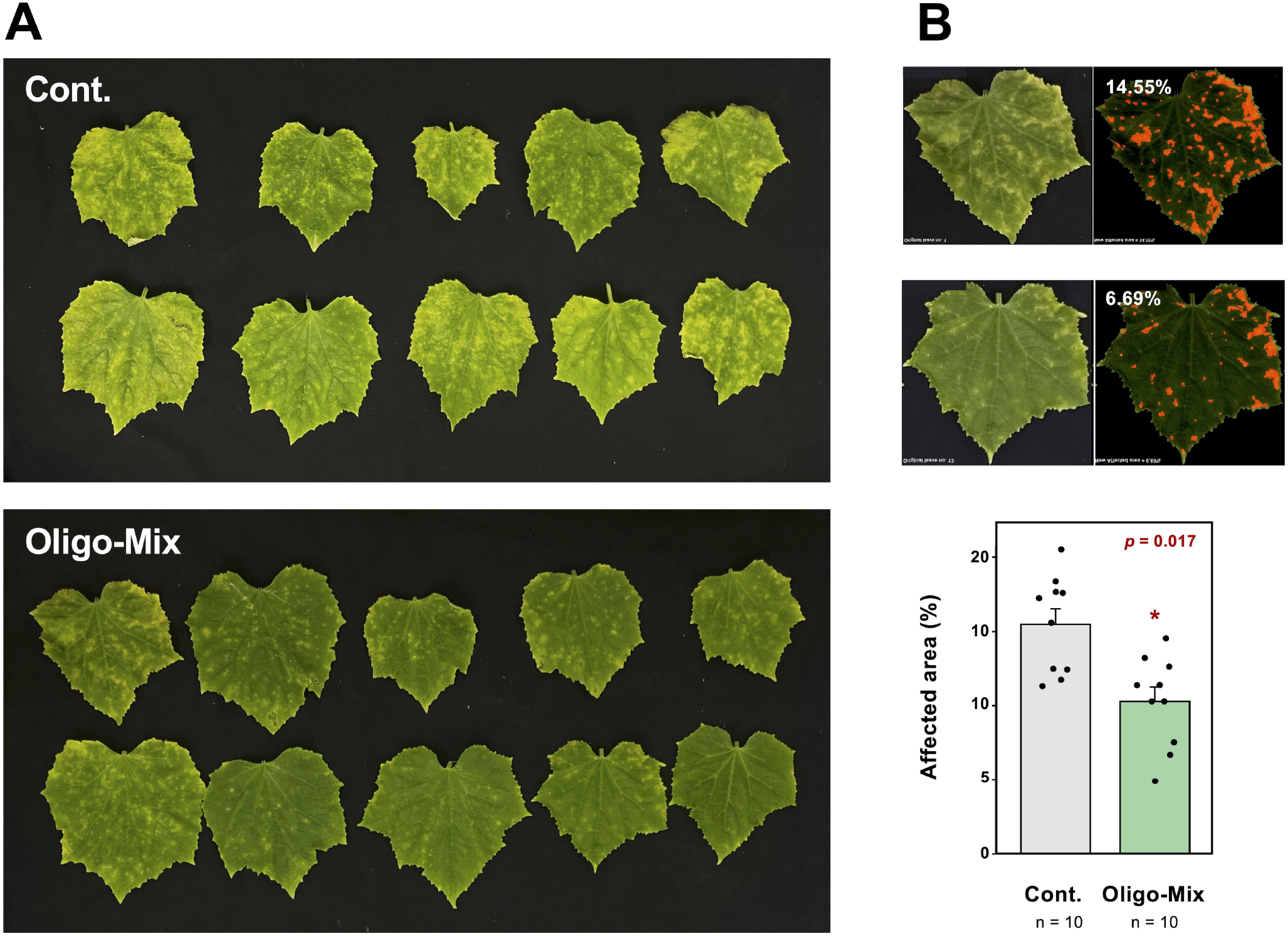
Oligo-Mix induces disease resistance of cucumber against anthracnose disease by *Colletotrichum orbiculare*. **(A)** Cucumber plants (4-weeks-old) were treated with control solution (Cont.) or Oligo-Mix (containing 20 µg/ml cello-oligosaccharide, 40 µg/m1 xylo-oligosaccharide and 20 µg/ml chitin-oligosaccharide) and spray-inoculated with spore suspension of *C. orbiculare* (1×10^5^ spores/ml, 0.5 ml/plant) at 24 h after treatment. Photographed 7 days after inoculation. **(B)** The diseased area of leaves was detected using an image analysis algorithm (See the method section) (n = 10). Asterisks indicate a significant difference from the control as assessed by a two-tailed Student’s *t*-test. * *p* < 0.05.

**Fig. 6.**
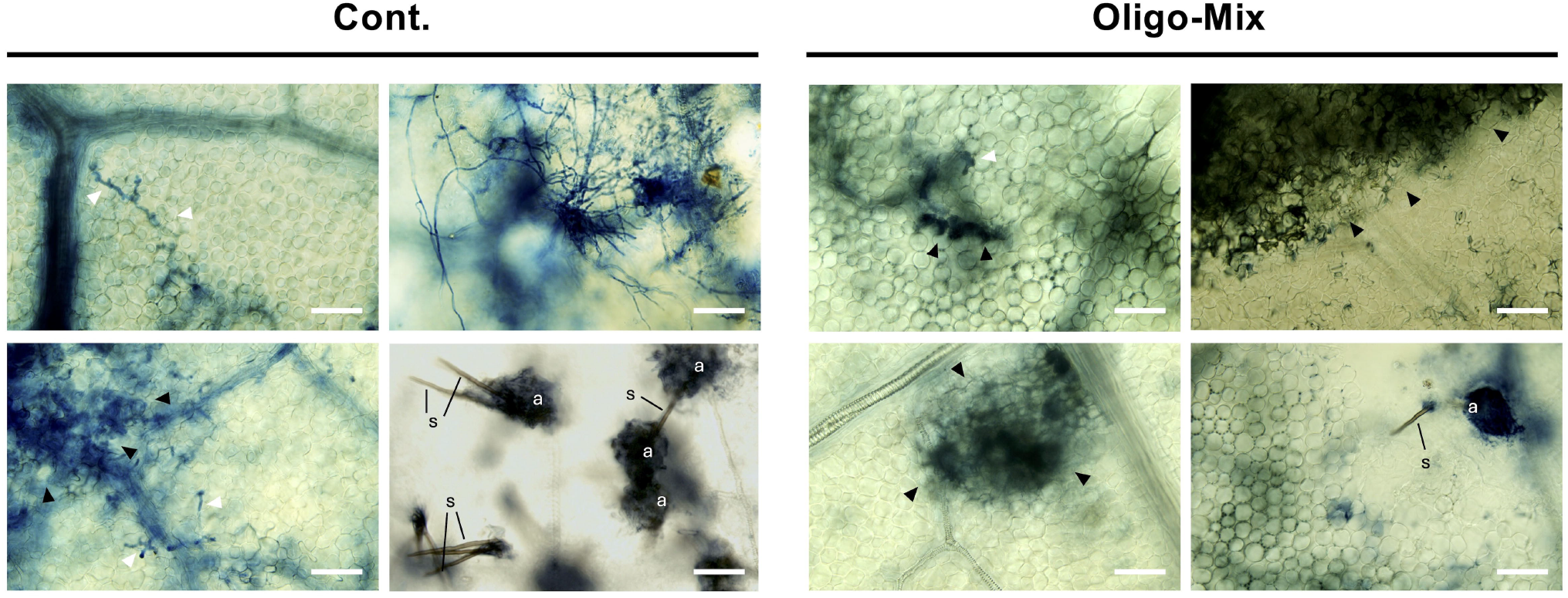
Microscopic observation of anthracnose infection on cucumber with or without Oligo-Mix treatment. Cucumber plants were treated with control solution (Cont.) or Oligo-Mix (20 µg/ml COS, 40 µg/ml XOS, 20 µg/ml CHOS) and conidial suspension of *Colletotrichum orbiculare* (1×10^5^ spores/ml, 0.5 ml/plant) were spray-inoculated. Inoculated leaves were stained with lacto-phenol trypan blue 8 days after the inoculation. In control cucumber leaves (left), intracellular biotrophic hyphae (white arrowheads) growing out from necrotic lesions (black arrowheads) and developing acervulus (a) with multiple setae (s) were often observed. In Oligo-Mix treated leaves, biotrophic hyphae were hardly detected and fewer and less-developed acervulus were observed. Bars = 50 µm.

### 3.4 Expression profile of cucumber genes in response to Oligo-Mix treatment

Four-weeks old cucumber leaves were treated with the control solution or Oligo-Mix, and expression of cucumber genes at 24 h after treatment was investigated by RNA-seq analysis. The list of significantly up-regulated genes (based on the average TPM score) in cucumber leaf is shown as Table S1. Oligo-Mix treatment upregulated 123 genes (FC > 2, p<0.05, average TPM>1) and downregulated 46 genes (FC < 0.5, p<0.05, average TPM>1). The upregulated genes include some genes predicted to be involved in cell reinforcement and expansion (Fig. 7A, Table S1). Exordium-like 1 (CsaV3_3G047580) encodes extracellular protein EXO mediates leaf cell expansion (Schröder et al., 2009). Xyloglucan endotransglucosylases/hydrolases (CsaV3_5G031640 and CsaV3_6G038030) are known as enzymes for the modification of cell wall structure by cleaving and re-joining xyloglucan in primary cell walls (Eklöf et al., 2010). Arabinogalactans (CsaV3_7G003940, CsaV3_4G009610 and CsaV3_7G028250) are a class of glycoproteins anchored to the plasma membrane, presumed to be involved in cellulose synthesis and deposition during plant cell wall biogenesis (Lin et al., 2022). Expression of genes for stress responses were also upregulated such as ERF/AP2 family transcription factors (e.g. CsaV3_3G016760, CsaV3_3G018600 and CsaV3_5G005890, Imano et al., 2022) and chaperon HSP70 (CsaV3_5G026520) (Fig. 7B, Table S1). Oligo-Mix treatment enhanced genes for plant disease resistance, including Avr9/Cf-9 rapidly elicited protein-like (CsaV3_4G025220, Rowland et al., 2005), F-box family protein (CsaV3_6G048900), CCR4-associated factor 1 homolog (CsaV3_6G038490, Liang et al., 2009), PAR1 protein (CsaV3_3G017150, Herbers et al., 1995, Takemoto et al., 2003) and thaumatin-like pathogenesis-related protein 5 (PR-5, CsaV3_3G045940, de Jesús-Pires et al., 2020). To determine the overall influence of treatment with Oligo-Mix on cucumber, Arabidopsis homologues of upregulated cucumber genes (Table S1) were assigned to gene ontology (GO) terms using the analysis tool PANTHER (Thomas et al., 2022) for GO enrichment analysis (Fig. S1). This analysis confirmed that Oligo-Mix can enhance the biological processes (BP) categories related to “defense response to fungus (GO:0050832)” and “defense response (GO:0006952)”. GO term “response to hormone (GO:0009725)” was also listed, but 11 genes under this category include genes for transcription factors predicted to be involved in the response to various plant hormones such as ethylene, abscisic acid and jasmonic acid, thus activation of specific hormonal responses by Oligo-Mix treatment was not identified.

**Fig. 7.**
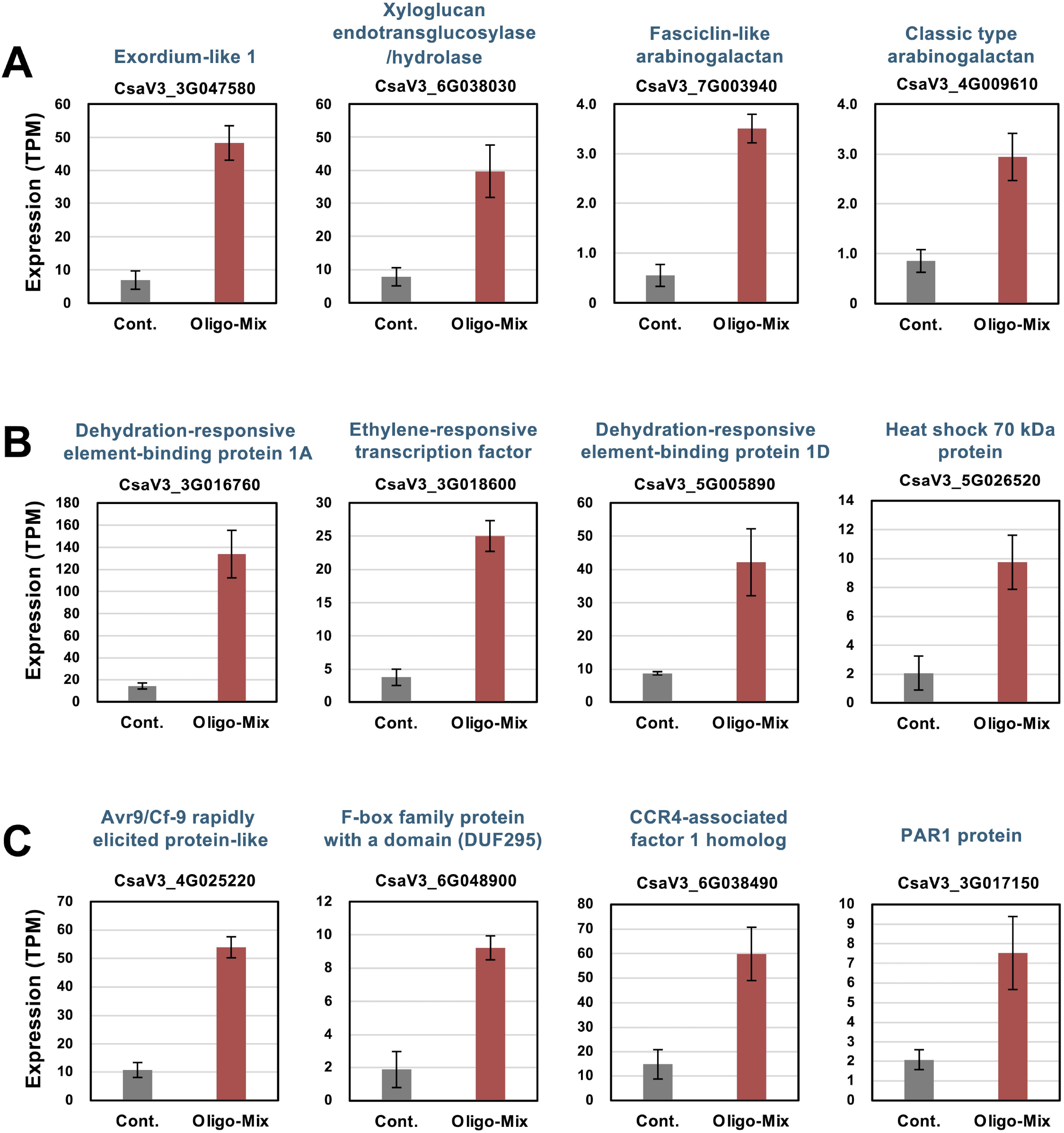
Expression profiles of representative cucumber genes upregulated by treatment with Oligo-Mix. Gene expression (TPM value) was determined by RNA-seq analysis of cucumber leaves treated with control solution (Cont.) or Oligo-Mix (20 µg/ml COS, 40 µg/ml XOS, 20 µg/ml CHOS) for 24 h. **(A)** Genes related to cell wall reinforcement. **(B)** Genes related to the responses to abiotic stresses. **(C)** Genes related to plant defense. Data marked with asterisks are significantly upregulated from control as determined by two-tailed Student’s *t-*test ** *p* <0.01, * *p* <0.05. See Table S1 for the list of upregulated genes by Oligo-Mix treatment.

The activation of genes associated with stress tolerance and disease resistance often comes at a trade-off with the plant growth, such as photosynthesis (Karasov et al., 2017; He et al., 2022; Godínez-Mendoza et al., 2023; Gao et al., 2024). We assessed the effect of Oligo-Mix treatment on highly expressed genes, however, Oligo-Mix treatment did not decrease the expression of genes involved in photosynthesis, but rather increased the expression of chlorophyll a-b binding protein, although this effect was not statistically significant (Table S2). These results indicate that Oligo-Mix treatment promotes the activation of cell growth and disease stress tolerance while maintaining the expression levels of genes required for photosynthesis.

### 3.5 Effect of Oligo-Mix treatment on powdery mildew resistance in greenhouse

The effect of Oligo-Mix treatment on disease incidence in cucumbers grown in greenhouses was investigated. Two-week-old cucumber seedlings grown in a controlled growth chamber were transferred to pots and grown in a greenhouse. Cucumber plants were sprayed with control solution or Oligo-Mix once every week (6 times altogether) and the number of naturally occurring colonies of powdery mildew on leaves was counted. Although no statistically significant differences were found because some plants did not show natural disease symptoms in both treatments, but the Oligo-Mix treatment tended to reduce the incidence of powdery mildew on cucumbers grown in the greenhouse (Fig. 8), suggesting the potential effectiveness of the practical use of Oligo-Mix in cucumber cultivation.

**Fig. 8.**
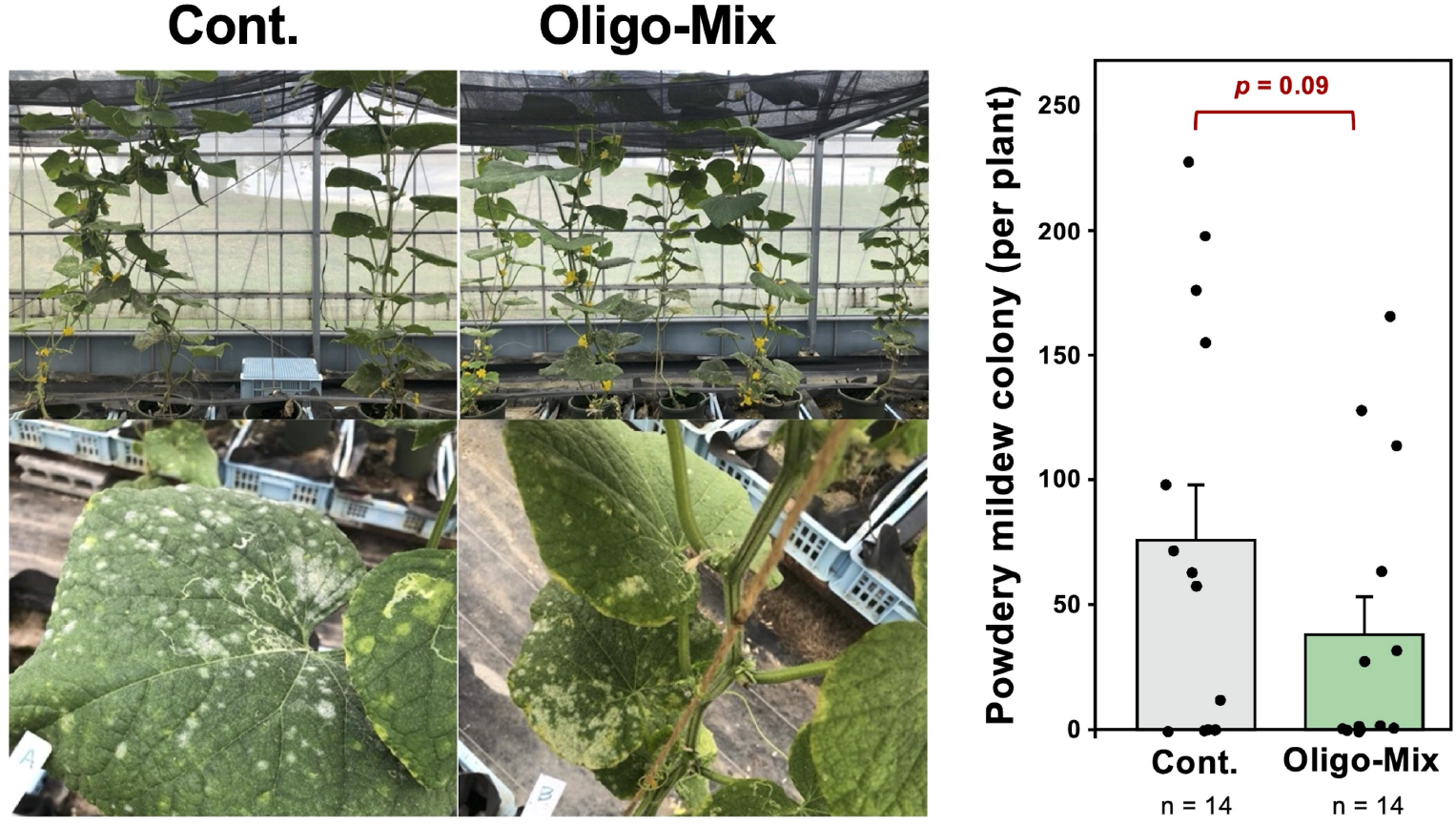
Effect of Oligo-Mix treatment on the appearance of powdery mildew on cucumber grown in greenhouse. Cucmber plant was sprayed control solution (Cont.) or Oligo-Mix (20 µg/ml COS, 40 µg/ml XOS, 20 µg/ml CHOS) once every week (6 times altogether) and naturally allowed to get infected by powdery mildew. The number of colonies per plant was counted at 6 weeks after first treatment. Data are shown as mean ± SD (n = 14). *p* value shown was calculated by a two-tailed Student’s *t*-test.

### 3.6 Conclusion remarks

The study suggests that treatment with Oligo-Mix, a mixture of the DAMPs COS and XOS and the MAMP CHOS, promotes both growth and disease resistance in cucumbers. Against the rational trade-off theory, why Oligo-Mix did not suppress the growth of plants while improving disease resistance? First, Oligo-Mix promoted root growth, which is expected to have the effect of allowing plants to grow basically robust and underpin the energy that is spent in the immune response. Second, the RNA-seq data indicated that the induced genes included some transcriptional regulators, and GO analysis identified activated molecular function (MF) are “Transcription cis-regulatory region binding (GO:0000976)” and “DNA-binding transcription factor activity (GO:0003700)”(Table S1, Fig. S1). Increased amounts of various transcriptional regulators are expected to allow plants to respond rapidly and robustly to the next environmental stresses (Van der Ent et al., 2009; Byun et al., 2014; Khan et al., 2022). Such effect is called priming, and the disease resistance induced by the Oligo-Mix might be at the level of priming (Conrath et al., 2002; Mauch-Mani et al, 2017; Hönig et al., 2023). Induction of priming is presumed to be less energetic loss than actual resistance to a pathogen that exhibits a resistance effect (e.g., accumulation of antimicrobial substances). In fact, hypersensitive cell death was not induced by Oligo-Mix treatment alone but was induced after inoculation with the powdery mildew (Fig. 4). Another possible explanation is that concentration of Oligo-Mix used in this study was optimal for inducing “eustress” of cucumber in the context of the hormesis model. Godínez-Mendoza et al (2024) suggested that in the eustress zone of the hermetic curve is characterized by the stimulation of growth and productivity and simultaneous activation of defense mechanisms.

A mixture of substances promoting growth and weakly activating disease resistance would be ideal combination as agricultural materials. Biostimulant materials, however, require further research and study to accumulate experiences for their effective use in agriculture, since different crops (and different cultivars) often have variable sensitivities to such substances.

## Acknowledgment

We are grateful to Ms. Yuki Iwamoto (Nagoya University) for her technical support. We also express our gratitude to Prof. Yoshitaka Takano (Kyoto University) for providing Colletotrichum orbiculare. We would like to acknowledge the Japanese Ministry of Education, Culture, Sports, and Technology (MEXT) for supporting Pring Sreynich in her pursuits. study in Japan on scholarship. This manuscript has been released as a pre-print on bioRxiv (Pring et al., 2024).

