## Supplemental Figure and Tables for "Mixed DAMP/MAMP oligosaccharides promote both growth and defense against fungal pathogens of cucumber"

Pring et al. 2025

**Table S1** List of cucumber genes significantly upregulated by treatment with Oligo-Mix (Fold change>2.5, p<0.05, average TPM>1).

| Gene ID | Cont. (Average TPM) | Oligo-Mix (Average TPM) | Fold change Oligo-Mix/Cont. | P value | Annotation | Homologous Arabidopsis gene | Gene ID | E-value |
|---|---|---|---|---|---|---|---|---|
| CsaV3_5G032180 | 0.00 | 2.86 | / | 0.03 | Pentatricopeptide repeat-containing protein | Not found | - | - |
| CsaV3_3G000800 | 0.14 | 2.20 | 15.44 | 0.03 | Chalcone-flavanone isomerase family protein | Chalcone-flavanone isomerase family protein | AT5G66230 | 1.0E-84 |
| CsaV3_6G031650 | 0.66 | 9.04 | 13.67 | 0.00 | Hypothetical protein | Not found | - | - |
| CsaV3_1G029100 | 0.69 | 6.66 | 9.64 | 0.04 | Hypothetical protein | Not found | - | - |
| CsaV3_3G016760 | 14.27 | 133.84 | 9.38 | 0.01 | Dehydration-responsive element-binding protein 1A | ERF/AP2 transcription factor family/C-repeat/DRE binding factor 2 | AT4G25470 | 2.0E-68 |
| CsaV3_3G047580 | 6.93 | 48.28 | 6.97 | 0.00 | EXORDIUM-like 1 | Phosphate-responsive 1 family protein | AT4G08950 | 6.0E-114 |
| CsaV3_3G016770 | 0.78 | 5.40 | 6.90 | 0.00 | Ethylene-responsive transcription factor | Ethylene-responsive transcription factor ERF025 | AT5G52020 | 1.0E-33 |
| CsaV3_5G031640 | 0.28 | 1.94 | 6.85 | 0.02 | Xyloglucan endotransglucosylase/hydrolase | Xyloglucan:xyloglucosyl transferase 33 | AT1G10550 | 6.0E-133 |
| CsaV3_3G018600 | 3.75 | 25.02 | 6.67 | 0.00 | Ethylene-responsive transcription factor | Ethylene-responsive transcription factor ERF109 | AT4G34410 | 2.0E-28 |
| CsaV3_7G003940 | 0.55 | 3.51 | 6.37 | 0.00 | Fasciclin-like arabinogalactan protein | FASCICLIN-like arabinogalactan protein 21 precursor | AT5G06920 | 4.0E-82 |
| CsaV3_1G029110 | 4.77 | 26.77 | 5.62 | 0.00 | Hypothetical protein | Not found | - | - |
| CsaV3_1G010540 | 0.35 | 1.84 | 5.31 | 0.03 | Hypothetical protein | Not found | - | - |
| CsaV3_6G038030 | 7.83 | 39.68 | 5.07 | 0.02 | Xyloglucan endotransglucosylase/hydrolase | Xyloglucan endotransglucosylase/hydrolase 15 | AT4G14130 | 9.0E-119 |
| CsaV3_4G025220 | 10.71 | 53.97 | 5.04 | 0.00 | Avr9/Cf-9 rapidly elicited protein-like | Protein of unknown function (DUF668) | AT3G23160 | 1.0E-79 |
| CsaV3_6G048900 | 1.90 | 9.23 | 4.86 | 0.01 | F-box family protein with a domain of unknown function (DUF295) | F-box protein with a domain protein | AT4G22030 | 5.0E-124 |
| CsaV3_5G005890 | 8.73 | 42.16 | 4.83 | 0.03 | Dehydration-responsive element-binding protein 1D-like | ERF/AP2 transcription factor family/DRE binding protein 1B | AT4G25490 | 7.0E-69 |
| CsaV3_5G026520 | 2.07 | 9.75 | 4.70 | 0.03 | Heat shock 70 kDa protein | Heat shock 70 kDa protein | AT5G02500 | 0 |
| CsaV3_4G036310 | 1.63 | 7.54 | 4.63 | 0.01 | GLUTAMINE DUMPER 5-like | GLUTAMINE DUMPER 4 | AT2G24762 | 2.0E-25 |
| CsaV3_2G032400 | 0.78 | 3.60 | 4.59 | 0.05 | Zinc finger protein 3-like | C2H2 and C2HC zinc fingers superfamily protein | AT5G10970 | 8.0E-35 |
| CsaV3_1G000650 | 5.46 | 24.28 | 4.44 | 0.04 | EF hand calcium-binding family protein | Calmodulin-like 38 | AT1G76650 | 7.0E-30 |
| CsaV3_6G031640 | 6.22 | 27.49 | 4.42 | 0.00 | Protein of unknown function (DUF241) | DUF241 domain protein | AT1G76240 | 2.0E-62 |
| CsaV3_5G006130 | 0.91 | 3.79 | 4.18 | 0.01 | Pentatricopeptide repeat-containing protein | Not found | - | - |
| CsaV3_1G012520 | 8.33 | 34.50 | 4.14 | 0.00 | Ethylene-responsive transcription factor | Ethylene-responsive transcription factor 12 | AT1G28360 | 2.0E-17 |
| CsaV3_6G050550 | 0.58 | 2.36 | 4.04 | 0.02 | Rac-like GTP-binding protein | Rac-like GTP-binding protein ARAC2 | AT5G45970 | 7.0E-120 |
| CsaV3_6G038490 | 14.82 | 59.86 | 4.04 | 0.02 | Polynucleotidyl transferase, ribonuclease H-like superfamily | CCR4-associated factor 1 homolog 11 | AT5G22250 | 4.0E-125 |
| CsaV3_7G013770 | 1.32 | 5.23 | 3.97 | 0.05 | Cysteine proteinase inhibitor | Cysteine proteinase inhibitor 5 | AT5G47550 | 3.0E-14 |
| CsaV3_3G007770 | 0.42 | 1.61 | 3.84 | 0.01 | Cytochrome b5-like | Cytochrome b5 isoform E | AT5G53560 | 2.0E-43 |
| CsaV3_5G014640 | 2.49 | 9.38 | 3.77 | 0.00 | Protein of unknown function, DUF538 | Protein of unknown function, DUF538 | AT1G09310 | 1.0E-54 |
| CsaV3_4G028360 | 4.58 | 16.95 | 3.70 | 0.02 | Ethylene-responsive transcription factor | Ethylene-responsive transcription factor ERF012 | AT1G21910 | 1.0E-30 |
| CsaV3_6G050310 | 4.31 | 15.94 | 3.70 | 0.03 | Hypothetical protein | Not found | - | - |
| CsaV3_3G017150 | 2.09 | 7.53 | 3.60 | 0.05 | PAR1 protein | Photoassimilate-responsive protein PAR-like | AT5G52390 | 2.0E-68 |
| CsaV3_5G023010 | 56.04 | 201.13 | 3.59 | 0.00 | AP2/ERF and B3 domain-containing transcription factor RAV1-like | AP2/ERF and B3 domain-containing transcription factor RAV1 | AT1G13260 | 5.0E-122 |
| CsaV3_6G002250 | 15.18 | 53.94 | 3.55 | 0.02 | Hypothetical protein | Not found | - | - |
| CsaV3_4G037790 | 9.78 | 34.58 | 3.54 | 0.01 | DUF4228 domain-containing protein | poly polymerase like | AT1G21010 | 4.0E-41 |
| CsaV3_4G009610 | 0.85 | 2.94 | 3.45 | 0.02 | Classical arabinogalactan protein 1-like | Not found | - | - |
| CsaV3_6G042660 | 2.00 | 6.80 | 3.40 | 0.00 | HTH-type transcriptional regulator protein ptxE | HTH-type transcriptional regulator | AT2G23690 | 8.0E-59 |
| CsaV3_6G038570 | 1.36 | 4.47 | 3.29 | 0.05 | RING-type E3 ubiquitin transferase | E3 ubiquitin-protein ligase PUB23 | AT2G35930 | 1.0E-134 |
| CsaV3_1G040660 | 118.24 | 389.42 | 3.29 | 0.00 | DDE Tnp4 domain-containing protein | DDE superfamily endonuclease | AT5G12010 | 0 |
| CsaV3_4G034970 | 156.81 | 509.31 | 3.25 | 0.01 | Zinc finger protein ZAT10-like | Zinc finger protein ZAT10 | AT1G27730 | 3.0E-58 |
| CsaV3_3G016800 | 0.48 | 1.53 | 3.17 | 0.02 | DETOXIFICATION protein | Multidrug and toxic compound extrusion protein 49 | AT4G23030 | 0 |
| CsaV3_3G040520 | 4.29 | 13.55 | 3.16 | 0.00 | Hypothetical protein | Not found | - | - |
| CsaV3_7G005810 | 50.21 | 155.69 | 3.10 | 0.00 | Ethylene-responsive transcription factor | Ethylene-responsive transcription factor 9 | AT5G44210 | 2.0E-29 |
| CsaV3_6G049930 | 1.27 | 3.92 | 3.09 | 0.01 | Dof zinc finger protein DOF1.4-like | Dof zinc finger protein DOF1.4 | AT1G28310 | 4.0E-23 |
| CsaV3_2G032650 | 1.40 | 4.22 | 3.02 | 0.03 | protein GLUTAMINE DUMPER 3-like | Glutamine dumper 5 | AT5G24920 | 9.0E-15 |
| CsaV3_7G003930 | 3.83 | 11.48 | 3.00 | 0.02 | Hypothetical protein | Hypothetical protein | AT3G09280 | 8.0E-09 |
| CsaV3_3G039370 | 153.88 | 445.35 | 2.89 | 0.01 | Hypothetical protein | Hypothetical protein | AT3G57450 | 4.0E-13 |
| CsaV3_5G028470 | 16.53 | 47.73 | 2.89 | 0.02 | Hypothetical protein | Hypothetical protein | AT3G02620 | 6.0E-35 |
| CsaV3_3G034460 | 0.95 | 2.73 | 2.88 | 0.01 | E3 ubiquitin-protein ligase | E3 ubiquitin-protein ligase RLIM-like protein | AT5G02020 | 5.0E-23 |
| CsaV3_7G028250 | 8.72 | 24.86 | 2.85 | 0.02 | classical arabinogalactan protein 26 | Not found | - | - |
| CsaV3_2G026210 | 162.89 | 464.10 | 2.85 | 0.01 | Zinc finger protein ZAT10-like | Putative salt-tolerance zinc finger protein | AT1G27730 | 7.0E-52 |
| CsaV3_6G047080 | 0.85 | 2.42 | 2.84 | 0.05 | WEB family protein | WEB family protein (DUF827) | AT3G51220 | 2.0E-27 |
| CsaV3_4G036890 | 24.03 | 67.01 | 2.79 | 0.01 | Hypothetical protein | Hypothetical protein | AT5G10750 | 1.0E-12 |
| CsaV3_2G034810 | 0.69 | 1.92 | 2.78 | 0.04 | Mitogen-activated protein kinase kinase kinase 2-like | Mitogen-activated protein kinase kinase kinase 17 | AT2G32510 | 3.0E-110 |
| CsaV3_6G003280 | 4.52 | 12.48 | 2.76 | 0.03 | Transmembrane protein | Hypothetical protein | AT1G62490 | 5.0E-18 |
| CsaV3_4G034040 | 3.16 | 8.72 | 2.76 | 0.01 | Transcription factor MYB44-like | Myb-related protein 77 | AT3G50060 | 6.0E-49 |
| CsaV3_5G028850 | 1.67 | 4.54 | 2.72 | 0.05 | Core-2/I-branching beta-1,6-N-acetylglucosaminyltransferase | Core-2/I-branching beta-1,6-N-acetylglucosaminyltransferase | AT1G68390 | 3.0E-136 |
| CsaV3_3G027820 | 4.42 | 12.03 | 2.72 | 0.00 | UPF0496 protein | Transmembrane protein, putative (DUF677) | AT3G19330 | 1.0E-42 |
| CsaV3_UNG000010 | 2.36 | 6.36 | 2.69 | 0.03 | Major latex-like protein | MLP-like protein 423 | AT1G24020 | 3.0E-03 |
| CsaV3_7G024550 | 9.25 | 24.91 | 2.69 | 0.00 | UPF0496 protein 4 | Hypothetical protein | AT1G19670 | 3.0E-142 |
| CsaV3_1G005530 | 1.79 | 4.82 | 2.69 | 0.03 | Disulfide bond formation protein B 2, putative | Hypothetical protein | AT3G47510 | 4.0E-11 |
| CsaV3_1G043180 | 24.60 | 65.87 | 2.68 | 0.01 | F-box protein | F-box family protein | AT1G22220 | 1.0E-38 |
| CsaV3_7G029910 | 10.79 | 28.73 | 2.66 | 0.03 | DUF4228 domain-containing protein | Hypothetical protein | AT3G03280 | 2.0E-06 |
| CsaV3_6G001690 | 119.81 | 318.19 | 2.66 | 0.04 | Nematode resistance protein-like HSPRO2 | Nematode resistance protein-like HSPRO2 | AT2G40000 | 2.0E-165 |
| CsaV3_5G025440 | 8.51 | 22.36 | 2.63 | 0.04 | ZnMc domain-containing protein | Matrix metalloproteinase | AT1G70170 | 1.0E-118 |
| CsaV3_4G033470 | 100.34 | 261.98 | 2.61 | 0.04 | Ethylene-responsive transcription factor 4 | Ethylene responsive element binding factor 4 | AT3G15210 | 2.0E-20 |
| CsaV3_2G035220 | 2.12 | 5.54 | 2.61 | 0.05 | Zinc finger protein CONSTANS-LIKE 12 | B-box type zinc finger protein with CCT domain | AT3G21880 | 4.0E-37 |
| CsaV3_6G036510 | 56.24 | 146.19 | 2.60 | 0.01 | GATA transcription factor | GATA transcription factor 8 | AT3G54810 | 1.0E-40 |
| CsaV3_5G025490 | 0.63 | 1.64 | 2.59 | 0.00 | RING-type domain-containing protein | RING/U-box superfamily protein | AT1G69330 | 2.0E-61 |
| CsaV3_7G028780 | 11.95 | 30.53 | 2.55 | 0.01 | Protein no-on-transient A-like | Not found | - | - |
| CsaV3_3G032400 | 7.64 | 19.34 | 2.53 | 0.00 | O-fucosyltransferase family protein | O-fucosyltransferase family protein | AT2G44500 | 0 |
| CsaV3_3G020040 | 3.41 | 8.57 | 2.52 | 0.03 | Unknown protein | Not found | - | - |
| CsaV3_2G004850 | 2.80 | 7.03 | 2.51 | 0.01 | TMV resistance protein N-like | Not found | - | - |

of cucumber

**Table S2** Effect of Oligo-Mix treatment on top 30 highly expressed gens in control cucumber leaves.

| Gene ID | Cont. (Average TPM) | Oligo-Mix (Average TPM) | Fold change | P value | Annotation |
|---|---|---|---|---|---|
| CsaV3_5G034360 | 67306.2 | 64013.0 | 0.95 | 0.6 | Ribulose bisphosphate carboxylase small subunit |
| CsaV3_5G008270 | 13790.7 | 14251.3 | 1.03 | 0.9 | Ribulose bisphosphate carboxylase/oxygenase activase, chloroplastic |
| CsaV3_UNG208380 | 12873.4 | 5942.2 | 0.46 | 0.5 | Photosystem II protein D1 |
| **CsaV3_3G031580** | **10981.3** | **14267.1** | **1.30** | **0.5** | **Chlorophyll a-b binding protein, chloroplastic** |
| **CsaV3_6G051530** | **10767.5** | **16513.0** | **1.53** | **0.3** | **Chlorophyll a-b binding protein, chloroplastic** |
| CsaV3_3G042440 | 9994.7 | 8218.3 | 0.82 | 0.2 | Carbonic anhydrase |
| **CsaV3_5G039350** | **8812.0** | **10368.1** | **1.18** | **0.6** | **Chlorophyll a-b binding protein, chloroplastic** |
| CsaV3_1G015100 | 8514.9 | 9768.1 | 1.15 | 0.6 | Photosystem I subunit O |
| **CsaV3_1G043710** | **8463.5** | **9913.4** | **1.17** | **0.5** | **Chlorophyll a-b binding protein, chloroplastic** |
| CsaV3_6G000050 | 7799.7 | 6742.4 | 0.86 | 0.7 | Hexosyltransferase |
| **CsaV3_6G013800** | **7495.3** | **10470.1** | **1.40** | **0.4** | **Chlorophyll a-b binding protein, chloroplastic** |
| **CsaV3_6G051520** | **7345.0** | **9912.0** | **1.35** | **0.4** | **Chlorophyll a-b binding protein, chloroplastic** |
| **CsaV3_7G002620** | **7171.3** | **10231.4** | **1.43** | **0.3** | **Chlorophyll a-b binding protein, chloroplastic** |
| **CsaV3_4G026100** | **6273.7** | **7263.8** | **1.16** | **0.6** | **Chlorophyll a-b binding protein, chloroplastic** |
| CsaV3_2G015280 | 6226.8 | 7133.5 | 1.15 | 0.6 | Fructose-bisphosphate aldolase |
| CsaV3_6G042680 | 6091.0 | 5507.8 | 0.90 | 0.2 | Oxygen-evolving enhancer protein 1, chloroplastic |
| CsaV3_UNG196500 | 5993.8 | 4876.5 | 0.81 | 0.8 | ORF43m |
| CsaV3_1G041670 | 5948.0 | 6411.2 | 1.08 | 0.8 | Arabinogalactan peptide 13-like |
| CsaV3_UNG209010 | 5866.3 | 4530.5 | 0.77 | 0.8 | 50S ribosomal protein L2, chloroplastic |
| CsaV3_4G007600 | 4664.4 | 3735.4 | 0.80 | 0.2 | Photosystem II 10 kDa polypeptide, chloroplastic |
| CsaV3_UNG203700 | 4483.2 | 3366.7 | 0.75 | 0.7 | ATP synthase subunit beta, chloroplastic |
| CsaV3_4G010600 | 4396.0 | 3754.8 | 0.85 | 0.8 | 30S ribosomal protein S3, chloroplastic |
| CsaV3_UNG240540 | 4379.6 | 2097.9 | 0.48 | 0.3 | LOW QUALITY PROTEIN: uncharacterized protein LOC112489936 |
| CsaV3_3G019220 | 4363.7 | 4687.8 | 1.07 | 0.5 | Ferredoxin |
| **CsaV3_5G030270** | **4359.9** | **5274.1** | **1.21** | **0.6** | **Chlorophyll a-b binding protein, chloroplastic** |
| CsaV3_1G026750 | 4342.7 | 6079.9 | 1.40 | 0.4 | Bifunctional inhibitor/lipid-transfer/seed storage 2S albumin superfamily |
| CsaV3_7G034490 | 4276.9 | 4712.2 | 1.10 | 0.7 | Germin-like protein |
| CsaV3_3G020110 | 4233.4 | 4258.8 | 1.01 | 1.0 | Glyceraldehyde-3-phosphate dehydrogenase |
| CsaV3_1G040810 | 4005.2 | 4666.9 | 1.17 | 0.4 | Photosystem II PsbX |
| CsaV3_4G037480 | 3956.8 | 3948.3 | 1.00 | 1.0 | Glyceraldehyde-3-phosphate dehydrogenase |
| **CsaV3_1G032510** | **3892.7** | **5551.1** | **1.43** | **0.5** | **Chlorophyll a-b binding protein, chloroplastic** |

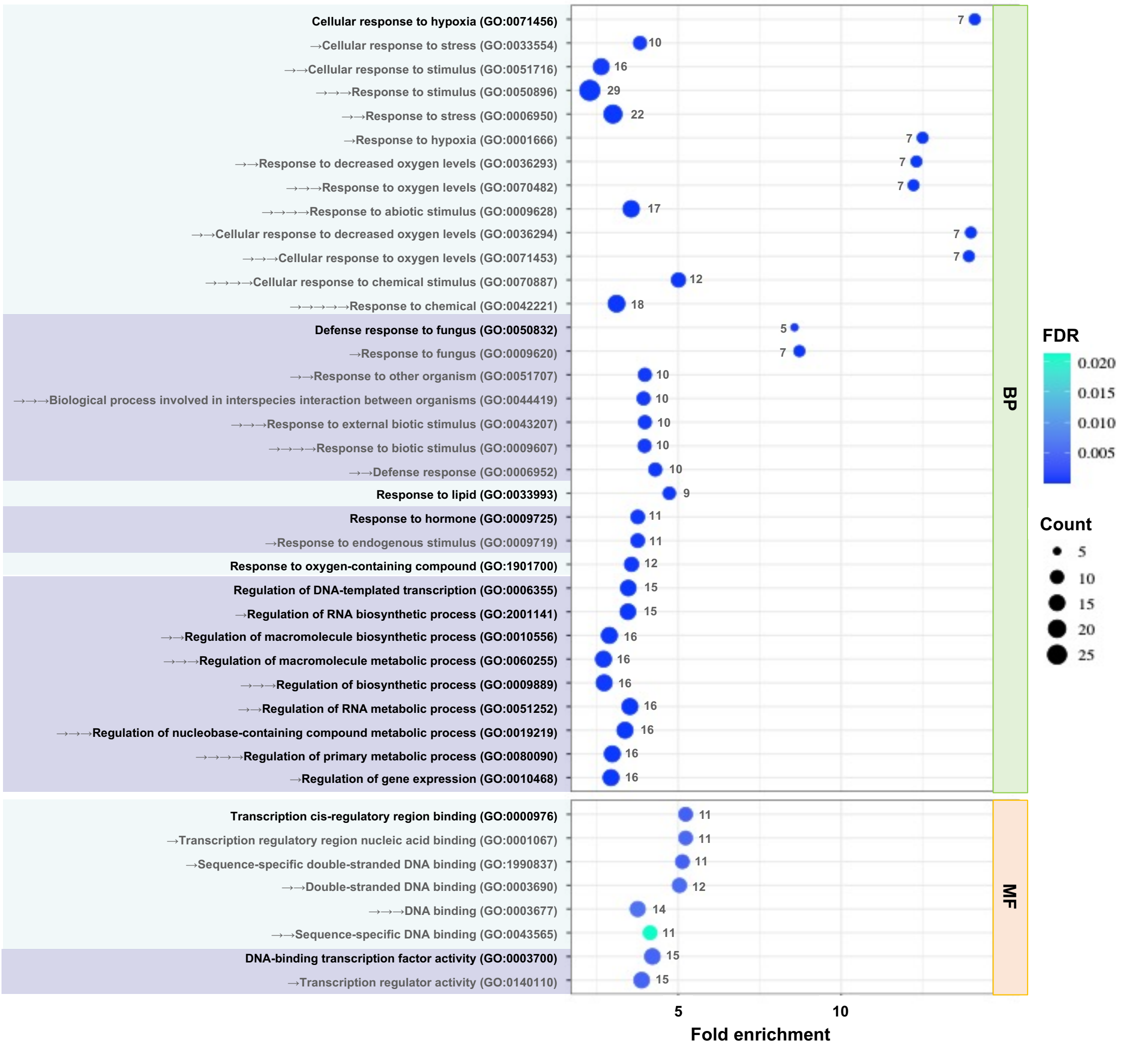


**Figure S1** Gene ontology (GO) enrichment analysis of up-regulated genes (Fold change > 2.5, $p < 0.05$) in cucumber treated with Oligo-Mix for 24 h. List of Arabidopsis homologous genes (See Table S1) were used for this analysis. GO term enrichment is expressed as significantly different fold enrichment of mapped genes (FDR $p<0.05$). The dot size (and numbers beside the dots) indicates the number of significantly up-regulated genes associated with the process and the dot color indicates the significance of the enrichment. BP, biological processes; MF, Molecular function.
